# FLIM-FRET as a Molecular Filter for Membrane-Induced Aggregation

**DOI:** 10.64898/2026.03.14.711702

**Authors:** Abdelrahman Salem, Wenzhu Qi, Jean-Christophe Rochet, Kevin J. Webb

**Affiliations:** Purdue University, West Lafayette, Indiana, 47907, USA

## Abstract

Membrane binding is thought to trigger early aggregation of alpha-synuclein (aSyn) in neurons. However, live-cell measurements of membrane-proximal aggregation with high specificity remain challenging. We combine three-channel fluorescence lifetime imaging microscopy (FLIM) with Förster resonance energy transfer (FRET) in a model that uses FRET as a proximity filter. A membrane-tethered donor reports membrane-aSyn separation through lifetime changes, while an aSyn-labeled acceptor reports the aggregation state through its lifetime. We estimate per-cell lifetimes and mixture fractions for membrane-bound and membrane-unbound populations using a hierarchical expectation-maximization (EM) algorithm that pools information across pixels. We validate the estimator using Monte Carlo studies. Using experimental neuronal data, this method resolves changes in membrane-proximal aggregation and aggregate-associated lifetimes. This framework provides quantitative per-cell metrics linking membrane proximity and aggregation for comparative live-cell studies.

**SIGNIFICANCE:** Many studies map FRET signals pixel by pixel or average signals over whole cells. Neither approach reliably answers key biological questions: how much membrane-proximal aggregation exists in a given cell, and how does it change across conditions? Here, we combine three-channel FLIM-FRET with a hierarchical analysis that estimates per-cell lifetimes and fractions for bound and unbound populations by pooling information across pixels. This method shifts the focus from noisy images to cell-level metrics that support comparisons across treatments, time points, or genotypes. Simulations and neuronal data show improved accuracy under realistic photon budgets and reveal membrane-proximal aggregation effects that were unattainable otherwise. This approach is broadly applicable and extends beyond alpha-synuclein.

## 1 INTRODUCTION

Cellular membranes are not passive boundaries. Their surfaces organize biochemical reactions by confining molecules, aligning reaction partners, and modulating local biochemical environments (1). Proteins that partition between cytosolic and membrane-proximal pools can therefore exhibit distinct encounter rates and assembly pathways depending on where they reside. Consistent with this principle, membrane localization can accelerate intermolecular association under membrane-confined conditions (2). For proteins whose function or dysfunction depends on membrane interactions, a central measurement problem is to determine how much of a given protein population is membrane-proximal and how much remains outside that membrane-proximal pool in intact cells.

In many relevant systems, proteins also populate multiple biochemical states, such as monomeric and aggregate-associated forms. In such cases, membrane proximity and assembly state become entangled in diffraction-limited measurements because several molecular classes can contribute photons to the same measurement voxel. Alpha-synuclein (aSyn) is a biologically important example of this broader problem because it partitions between membrane-associated and non-membrane-associated states (3, 4), and its membrane interactions are closely linked to its aggregation behavior (5, 6). We model aSyn using four classes defined by membrane association and assembly state, as illustrated in Fig. 1. Near the membrane, species from these classes can lie within tens of nanometers of each other and contribute photons to the same voxel, making it difficult to extract a quantitative, per-cell measure of membrane-proximal aggregation. Because conventional fluorescence microscopy integrates signals from these classes within a diffraction-limited volume, they are difficult to distinguish using intensity or just localization (7). In principle, super-resolution approaches such as STORM and PALM can improve spatial separation (8–10). However, in live-cell settings they still face practical tradeoffs in labeling, acquisition, and photon budget, and localization alone does not directly assign biochemical state to the emitting species (10).

**Figure 1:**
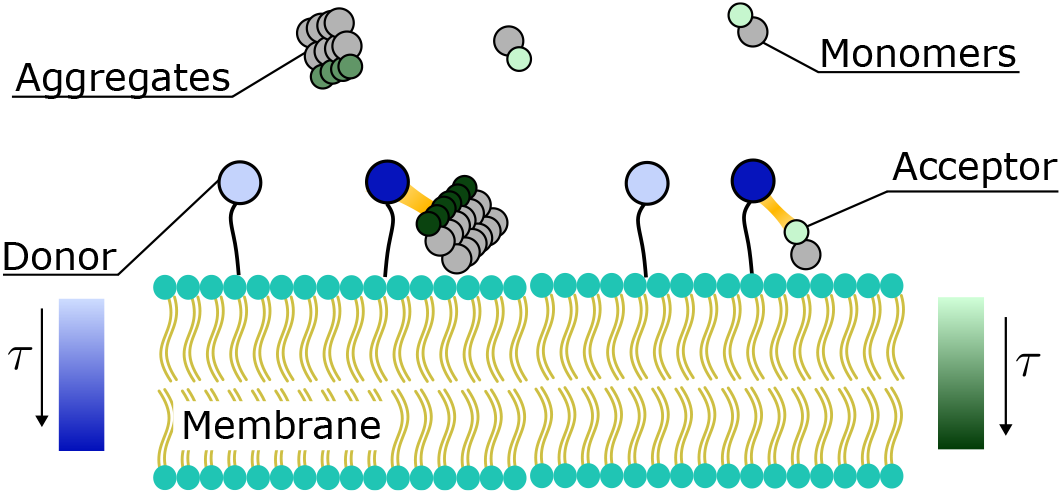
Conceptual schematic of the four aSyn molecular classes defined by membrane association (membrane-bound versus cytosolic) and assembly state (monomeric versus aggregate-associated). Donor fluorophores (blue) label the lipid bilayer membrane and report nanoscale membrane proximity of aSyn via donor-acceptor FRET, while acceptor self-quenching indicates aggregate-associated microenvironments. Donor-acceptor FRET and aggregation-associated quenching shorten the fluorescence lifetimes (*τ*) of the donor (left) and acceptor (right). Thus, donor lifetime reductions describe membrane proximity, whereas acceptor lifetime reductions show aggregation.

Fluorescence lifetime imaging (FLIM) provides a route to molecular-state information below the diffraction limit by measuring the lifetime – the time a fluorophore spends in its excited state – rather than where it is in space. The excited-state lifetime reflects local non-radiative de-excitation pathways – including FRET and quenching by neighboring molecules – and is therefore sensitive to changes in the microenvironment (11–13), while remaining largely independent of fluorophore concentration and excitation intensity. By introducing a suitable donor-acceptor pair, Förster resonance energy transfer (FRET) adds a well-defined, nanometer-scale distance sensitivity: donor lifetimes are shortened and acceptor emission is sensitized when donor and acceptor are within a few nanometers. In this work, donor-acceptor FRET reports nanometer-scale membrane proximity of labeled aSyn, while acceptor lifetime changes report aggregation-associated microenvironments. By combining these signatures in a joint model, we separate membrane-proximal and membrane-unbound pools that coexist within the same diffraction-limited voxel (14, 15), bypassing the need to optically resolve individual complexes (12, 16).

In practice, most cellular FLIM-FRET analyses rely on a single channel. Here, a channel denotes one excitation-emission configuration: a specific excitation wavelength combined with a defined emission band that yields one time-resolved decay curve per pixel. A lifetime decrease measured in a single channel can arise from multiple, indistinguishable causes, including proximity-dependent FRET (4), aggregation-associated quenching (12, 16), changes in local refractive index, or some mixture of these effects (13, 14). The underlying mixture decay is often a multi-exponential or non-exponential rather than a single exponential, which makes multi-parameter inference from a single-channel decay mathematically ill-posed. As a result, multiple parameter combinations can produce nearly identical curves, especially at low photon counts (17). Pixel-wise fitting further amplifies noise (18), while standard global fits (19–21) typically assume simpler decay structures and do not explicitly encode the four-class membrane-proximity and aggregation mixture considered here. Consequently, conventional single-channel FLIM-FRET analyses struggle to provide unique, stable, per-cell readouts of membrane-proximal aggregation.

This measurement challenge is biologically important for aSyn because its partitioning between membrane-bound and membrane-unbound states is closely linked to both its physiological function and its aggregation behavior. In neurons, a substantial fraction of aSyn interacts with phospholipid membranes (3, 4, 22), and these interactions are linked to its physiological role in synaptic vesicle trafficking and release (23, 24). At the same time, membrane binding can alter aSyn conformation and local concentration in ways that favor self-assembly at the membrane surface (3, 4, 23, 25). Bulk-phase nucleation is kinetically unfavorable at physiological concentrations (5, 26). In contrast, membrane binding can stimulate primary nucleation and co-aggregation (5, 27, 28). This makes aSyn a useful example of a broader class of membrane-mediated assembly problems in which the key unresolved quantity is not simply total aggregation, but the extent of membrane-proximal aggregation in individual cells.

This question is biologically important because aSyn aggregation is a hallmark of Parkinson’s disease and related synucleinopathies (29). Membrane-proximal aggregation has been proposed to contribute to early pathogenic events and to seed downstream spread between cells (30–34). Yet a remaining challenge is to obtain calibrated, per-cell readouts of membrane-proximal versus non-proximal aSyn populations together with their aggregation-associated lifetimes in intact neurons.

In this work, we develop and validate a three-channel FLIM-FRET framework for this problem. We treat FRET as a molecular filter for nanoscale membrane proximity and use acceptor self-quenching as a complementary signature of aggregate-associated microenvironments. In this setup, donor quenching and sensitized emission constrain membrane proximity, while acceptor lifetime shortening reports aggregation-associated states. Experimentally, we acquire three time-resolved measurements per field of view using distinct excitation and emission selections, which provide spectrally separated information to constrain donor quenching, sensitized acceptor emission, and acceptor lifetime changes. A forward model couples donor-acceptor proximity on the membrane to these measurements and makes spectral mixing, gain, and noise explicit. Rather than fitting each pixel independently, we formulate a hierarchical inference problem targeted to biologically meaningful cell-level quantities, including the lifetimes and fractions of the four aSyn classes in each cell. This approach pools information across pixels in a statistically principled manner and yields robust, low-variance per-cell readouts under experimental conditions.

We fit the model with an expectation-maximization (EM) algorithm that alternates between pixel-wise weighted updates under the known noise model and global updates of cell-level parameters that summarize the pixel distribution (35, 36). This formulation yields per-cell estimates with reduced variance at typical photon counts while preserving interpretability in terms of aggregate fraction, membrane-proximal fraction, and aggregate-associated lifetime changes. We validate the approach using a Monte Carlo method on datasets with known simulated values, comparing the proposed hierarchical EM framework against a conventional pixel-wise fit followed by cell averaging. We then apply the framework to neuronal measurements and quantify the per-cell differences obtained by the hierarchical analysis when extracting membrane-proximal aggregation readouts from the three-channel data. Together, these results provide a calibrated path from three-channel FLIM-FRET measurements to biologically specific, per-cell readouts of membrane-induced aggregation under realistic photon counts.

## 2 MATERIALS AND METHODS

### 2.1 Imaging platform and acquisition

We carried out time-gated fluorescence lifetime imaging on a custom-built wide-field FLIM system based on an Olympus iX73 inverted microscope as illustrated in Fig. 2(a). Fluorescence was excited through the objective and detected in a gated image-intensified CMOS camera operated in a time-gated mode. For each pixel, we acquired three FLIM channels corresponding to donor (D), sensitized emission (S), and acceptor (A) signals. We defined these channels by the excitation and emission filter combinations mounted in interchangeable filter cubes in the microscope turret (Fig. 2(b)). Channel D used donor excitation (430 nm) with donor-band emission detection (460–480 nm), Channel S used donor excitation (430 nm) with acceptor-band emission detection (500–550 nm) to isolate sensitized acceptor fluorescence, and Channel A used acceptor excitation (488 nm) with acceptor-band emission detection (500–550 nm). Throughout the manuscript, these three channels define the FLIM-FRET measurements.

**Figure 2:**
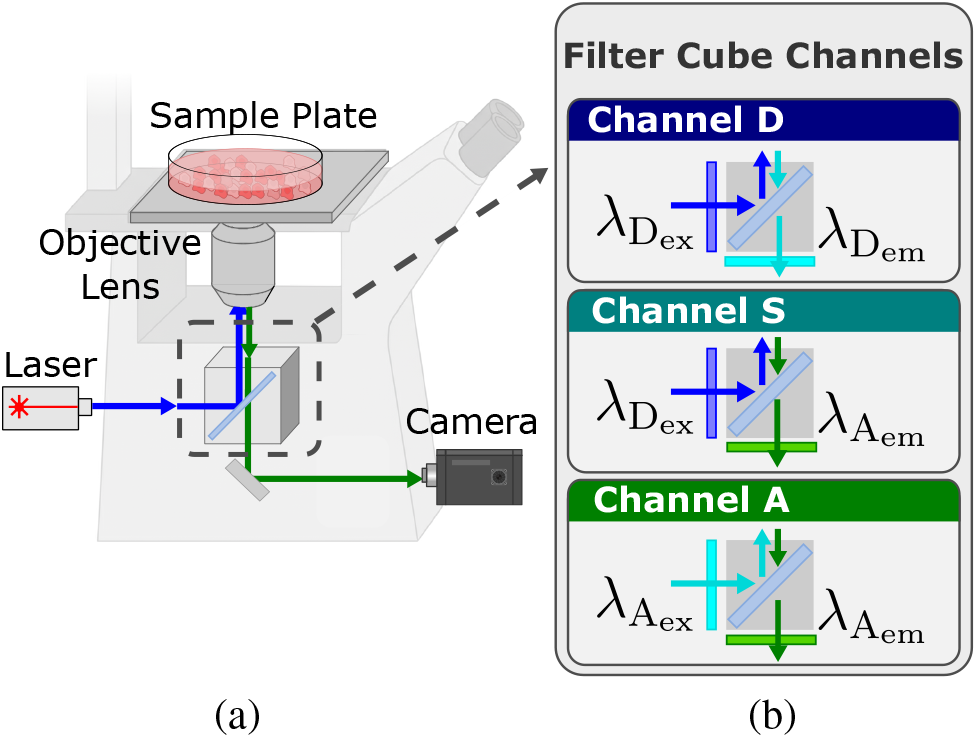
Schematic of the three-channel FLIM-FRET microscope and filter-cube configuration. (a) Epifluorescence microscope used for FLIM-FRET measurements. A laser source is delivered through the objective lens to illuminate the sample, and fluorescence is collected by the same objective and relayed to the camera through a filter cube. (b) Filter-cube configurations defining different channels by excitation/emission combinations. Channel D, the donor channel, uses donor-excitation 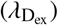 and donor-emission 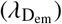 filters; Channel S, the sensitized acceptor emission channel, uses donor-excitation and acceptor-emission 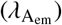 filters; and Channel A, the acceptor channel, uses acceptor-excitation 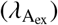 and acceptor-emission filters to record acceptor-only fluorescence. These three channels are used throughout the manuscript to define the measured FLIM-FRET signals.

The excitation source was a 1 MHz, 1040 nm femtosecond laser (Spirit, Spectra-Physics) coupled to an optical parametric amplifier (OPA), providing tunable excitation across 320–1040 nm with ~150 fs pulse duration. After the OPA, the beam power was attenuated to 1 mW at the sample, corresponding to a pulse energy of approximately 1 nJ for live-cell imaging, using a continuously variable neutral density filter (Thorlabs). We routed the laser beam into the microscope via mirrors and lenses to provide single-photon, wide-field epifluorescence illumination at the sample plane. We implemented time-resolved detection using a LaVision PicoStar HR12 image intensifier coupled to an Imager-M-Lite CMOS camera. A picosecond delay generator, synchronized to the laser trigger output, controlled the relative timing between excitation pulses and the intensifier gate. We sampled fluorescence decays in 200 ps increments with a 200 ps gate width and a 500 ms exposure time per gate, yielding 111 gates spanning a 22 ns acquisition window. The temporal instrument response function *h* (*t*) was measured using Allura Red dye with 10 ps lifetime (37) and normalized to unit area. A complete three-channel FLIM dataset required approximately 3 minutes of acquisition, corresponding to 2.4 ×10^8^ excitation pulses and a total delivered energy of about 240 mJ. For an illuminated area ≈0.25 mm^2^, this corresponds to a radiant exposure of 96 J cm^−2^. Under these conditions, we observed minimal photobleaching and no acute morphological changes during imaging. Consistent with established phototoxicity guidelines, irradiance and radiant exposure at the sample plane remained within a non-phototoxic regime, and neuronal cultures retained stable morphology and continued neurite outgrowth over more than 7 days of repeated imaging (38).

### 2.2 Forward model formulation

This section formulates a forward model that maps a small set of biophysical parameters to the measured three-channel time-resolved decays. We first derive the ideal channel decays *v*_D_(*t*), *v*_S_(*t*), and *v*_A_(*t*). Here, “ideal” refers to the decay profiles prior to measurement effects, that is, before instrument response, spectral bleed-through (emission leakage between detection bands), cross-excitation (off-target excitation due to spectral overlap), and additive background. We then assemble these effects into measurement operators and collect the full forward model in a compact matrix form.

To connect these three ideal decays through shared biophysical parameters, we model the aSyn population as four molecular classes defined by membrane association and assembly state: membrane-bound monomers (bm), membrane-unbound monomers (um), membrane-bound aggregates (ba), and membrane-unbound aggregates (ua). This biological parameterization is necessary because the kinetics of aSyn are fundamentally dictated by its physical environment and its degree of oligomerization. We collect their respective population fractions into the set

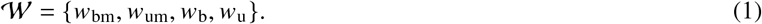

These fractions satisfy the probability simplex constraints

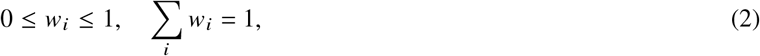

where *i* ∈ {bm,um,b,u} indicates the four molecular classes.

#### 2.2.1 Donor channel and membrane FRET

We derive the ideal decay, *v*_D_ (*t*), for the donor channel. Traditional FRET models assume a 1:1 donor-to-acceptor interaction. However, because membrane-bound donors interact with a two-dimensional field of multiple, randomly distributed acceptors in the lipid bilayer, we must account for this spatial distribution. Accordingly, we assume that these acceptors are independently distributed in the membrane plane with an effective surface density *ρ* (acceptors per unit membrane area), and a minimum donor-acceptor separation *R*_e_ such that donor-acceptor separation, *r*_da_, satisfy *r*_da_ ≥ *R*_e_, which accounts for the finite size of the fluorescent-protein tags (39–42). For example, GFP-like fluorescent proteins form a ~ 2.4 nm diameter *β*-barrel (43), with one barrel on each side of the donor-acceptor contact, this gives *R*_e_ = 4.8 nm (44). We also fix the Förster distance *R*_0_ = 5.83 nm (the donor-acceptor separation at which FRET efficiency equals 50%) from published photophysics (45) for the chosen fluorophore pair, and we fix the unquenched donor lifetime *τ*_d_ to values obtained from donor-only calibration samples, as described in the Methods Sect. 2.6. Under this model, we use the Wolber-Hudson expression to describe the ensemble average decay profile as (39)

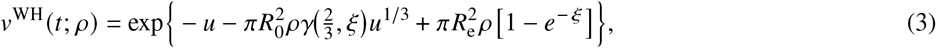

where *γ* is the lower incomplete gamma function, *u* = *t*/ *τ*_d_, and *ξ* = (*R*_0_ /*R*_e_)^6^*u*. This model assumes a laterally random in-plane acceptor distribution, so nonrandom clustering or out-of-plane geometry may introduce model mismatch. Accordingly, the inferred surface densities should be interpreted as effective parameters under such conditions (39–41).

In the membrane-bound population, we model two effective acceptor microenvironments that differ only in acceptor surface density: a monomer domain with *ρ* = *ρ*_m_ and an aggregate domain with *ρ* = *ρ*_a_. The ideal donor-channel decay profile is then a mixture over both domains, giving

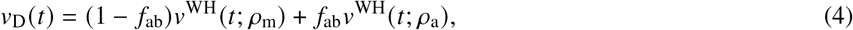

with mixture fraction *f*_ab_, which defines the fraction of aggregates among membrane-bound proteins and is derived from the acceptor-state mixture as

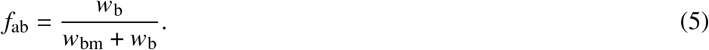

This choice ties donor domain occupancy to the composition of membrane-bound acceptors. For interpretability, we summarize the donor decay profile by an equivalent lifetime and FRET efficiency, which provide an intuitive measure of donor–acceptor proximity in this membrane model. The equivalent quenched donor lifetime is

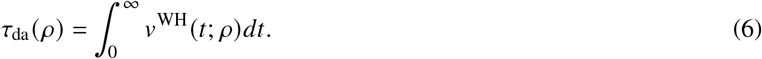

This corresponds to an equivalent FRET rate

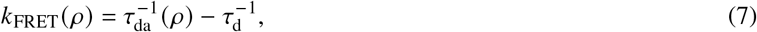

where *τ*_d_ is the unquenched donor lifetime. The equivalent FRET efficiency for a given acceptor density is

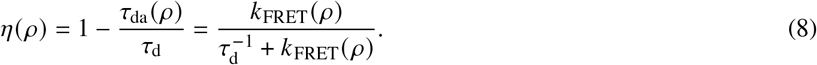

This corresponds to a total equivalent FRET efficiency of

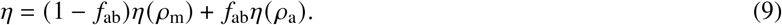

These expressions complete the donor-channel component by defining *v*_D_ *t* and its derived proximity summaries *τ*_da_ and *η* in terms of *ρ*_m_, *ρ*_a_, and *f*_ab_.

#### 2.2.2 Sensitized channel donor-to-acceptor excitation

To model this sensitized pathway, we approximate the complex, non-exponential donor decay in Eq. (3) as an effective first-order process with equivalent mono-exponential lifetime in Eq. (6). This yields the excited donor population, *N*_D_(*t*) ≈ *N*_D_(0) exp (− *t*/*τ*_da_(*ρ*)). This energy transfer acts as the driving source for the sensitized acceptor population, *N*_S_(*t*), whose dynamics are governed by the state equation

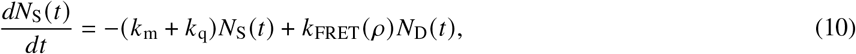

where *k*_m_ is the intrinsic monomer decay and *k*_q_ is a non-radiative self-quenching rate associated with aggregate microenvironments. As aggregation increases, the total de-excitation rate *k*_m_ + *k*_q_ becomes larger and the effective lifetime becomes shorter. The physical origin of *k*_q_ is concentration-dependent self-quenching of mVenus in aggregate microenvironments: at high local fluorophore densities, excited-state energy migrates via FRET to nonfluorescent trap sites formed by proximal fluorophore pairs (46), and fluorescence lifetime reductions consistent with this behavior have been directly observed for aggregating fluorescent-protein-tagged constructs (12, 47). In the membrane-bound aggregates, we define *k*_q_ = *k*_b_, whereas we define *k*_q_ = *k*_u_ for the membrane-unbound aggregates. These rates result in three distinct lifetimes

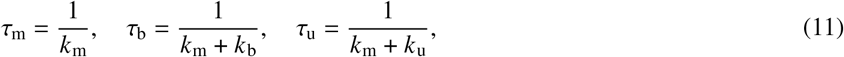

where *τ*_m_ corresponds to the monomers, whether membrane-bound (bm) or membrane-unbound (um). This definition for the lifetimes implicitly enforces the constraint

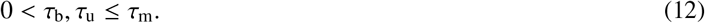

The solution of Eq. (10) has the form (48)

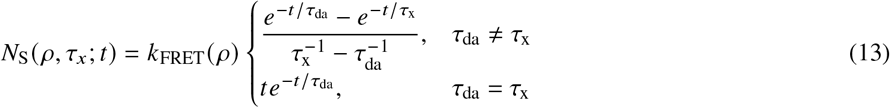

where *τ*_x_ represents either *τ*_m_, or *τ*_b_. The sensitized channel signal depends only on membrane-bound acceptor fractions, since unbound populations (*w*_um_, *w*_u_) are by definition distant from the donor-labeled membrane rendering their FRET efficiency negligible. The resulting ideal decay profile is

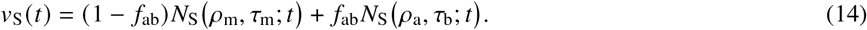

This completes the sensitized component by defining *v*_S_ (*t*) as donor-driven acceptor kinetics parameterized by *k*_FRET_ (*ρ*) and the acceptor lifetimes.

#### 2.2.3 Acceptor channel and aSyn aggregation

In the acceptor channel, we directly excite the acceptor population and record acceptor-band emission, so the kinetics reflect acceptor decay and aggregation-associated quenching without donor-driven excitation. We describe the excited-state population of acceptors, *N*_A_, using an assumed first-order state equation

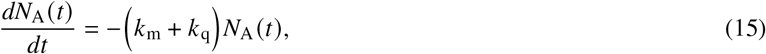

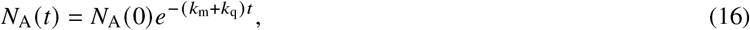

where *N*_A_ (0) is the initial acceptor population. We use the four molecular classes defined earlier with corresponding fractions in Eq. (1) and the three lifetimes defined in Eq. (11). The decay profile becomes

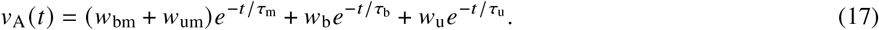

Thus, Channel A constrains aggregation through (*τ*_b_, *τ*_u_) and (*w*_ba_, *w*_ua_), independent of donor-driven excitation.

#### 2.2.4 Fluorescence decays measurement model and calibration operators

The preceding sections established the ideal theoretical photophysics of the fluorophores in isolation. To connect these idealized models to our experimental observables, we must now construct a macroscopic measurement model. This model uses the ideal decay profiles while accounting for the physical realities of the imaging system: finite temporal resolution, spectral cross-talk between detection channels, and detector noise. Given *v*_c_(*t*), where *c* ∈ {D,S,A}, as the ideal channel decay profiles defined in Eqs. (4), (14), and (17), define a time-sampled version 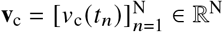. Let the instrument response function (IRF, *h*(*t*)) be sampled on 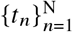. It is convenient to express convolution by a Toeplitz matrix **H** ∈ ℝ^N×N^ so that the convolution of the vector **v**_c_ ∈ ℝ^N^ with **H** is **Hv**_c_.

The measured decay in a given channel is generally a linear combination of the ideal decays *v*_D_ (*t*), *v*_S_ (*t*), and *v*_A_ (*t*) because of finite spectral separation in both excitation and emission. We therefore impose a physically structured mixing model determined by the optical signal paths. The diagonal elements are fixed to unity so that each channel is referenced to its primary ideal decay, while off-diagonal coupling arises through emission bleed-through between the donor and acceptor detection bands, parameterized by *α*_a_ and *α*_d_, and cross-excitation under the non-primary excitation wavelength, parameterized by *χ*_d_ and *χ*_a_. Under this factorized signal-path model, the spectral-mixing matrix **S** becomes

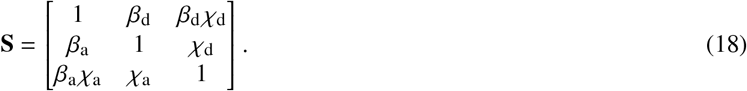

Furthermore, we scale the channel decay profiles by a scalar amplitude *α*_c_ describing the excitation power, absorption coefficient, detection emission, quantum yield, and initial number of excited states. We arrange these amplitudes into a vector *α* to represent the three channel amplitudes

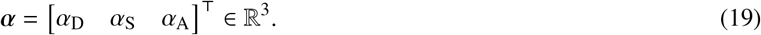

This model formulation results in the set of calibration parameters

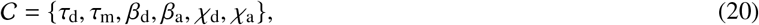

where *τ*_d_ and *τ*_m_ are the unquenched donor and acceptor lifetimes from donor-only and acceptor-only samples. The set 𝒞 is reserved for calibration constants estimated from dedicated control-sample experiments, namely donor-only and acceptor-only measurements as described in Methods Sect. 2.6; related calibration treatments are discussed in (48–50). Fixed geometric or photophysical quantities such as *R*_e_ and *R*_0_ are likewise prescribed independently rather than estimated from the control-sample calibration experiments. With 𝒞 fixed, the remaining unknowns are the biophysical parameters collected in the vector *θ* with seven degrees of freedom, resulting in

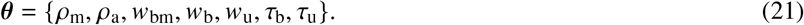

The remaining monomer fraction, *w*_um_, is strictly defined using the probability simplex as described by Eq. (2).

To isolate the non-linear physical parameters *θ*_*p*_ from the linear amplitudes *α* for channel *c*, we define **I**_*c,p*_ (*θ*_*p*_)^⊤^ as a row corresponding to the theoretical, unscaled, spectrally mixed decay matrix. Stacking these rows ordered (D,S,A), we construct the full forward model **I**_*p*_ (*θ*_*p*_, *α*_*p*_) ∈ ℝ^3×N^ by mimicking the physical signal path: the ideal, IRF-convolved decays are mapped through the spectral cross-talk matrix **S**, scaled by their respective amplitudes *α*_*p*_ via a diagonal matrix, and finally summed with the time-independent background **B** to give

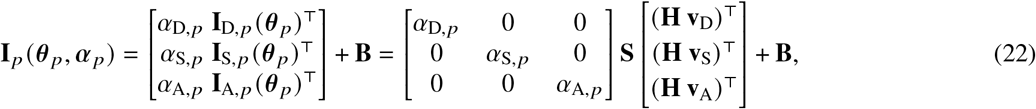

with **B** ∈ ℝ^3×N^ denoting the background matrix. Physically, **B** models time-uncorrelated photon counts, with each row taken to be constant across time bins. In our measurements, this background is dominated by the camera dark response, but it may also include other constant sources such as ambient stray light. For each acquisition setting, **B** is measured experimentally from camera frames acquired with no incident photons and is then held fixed during inference.

### 2.3 Biological parameters

The primary model parameters (*w*_bm_, *w*_um_, *w*_b_, *w*_u_) provide class-resolved fractions, but several biologically useful summaries are more naturally expressed as derived readouts. Accordingly, we define the aggregate, membrane-bound, and conditional membrane-associated fractions listed in Table 1 as deterministic functions of the inferred class fractions. These quantities are used throughout the Results section to summarize cell-level aggregation and membrane-proximity behavior in a more interpretable form.

**Table 1:** Definition of biological parameters extracted from experimental data.

| Variable | Description |
| --- | --- |
| $f_a = w_u + w_b$ | Fraction of aggregates among all proteins |
| $f_{ab} = \frac{w_b}{w_{bm} + w_b}$ | Fraction of aggregates among membrane-bound proteins |
| $f_b = w_{bm} + w_b$ | Fraction of membrane-bound proteins among all proteins |
| $f_{ba} = \frac{w_b}{w_u + w_b}$ | Fraction of membrane-bound aggregates among all aggregates |
| $f_{bm} = \frac{w_{bm}}{w_{um} + w_{bm}}$ | Fraction of membrane-bound monomers among all monomers |

### 2.4 Hierarchical cell-level estimation

Our goal is to obtain per-cell estimates of biologically interpretable parameters (lifetimes, fractions, and densities) rather than noisy per-pixel maps. To that end, we treat the three-channel forward model as a pixel-level likelihood and impose a simple hierarchical structure across pixels within each cell. Each pixel has its own physical parameter vector, but these parameters are coupled through an unconstrained latent representation in which we place a multivariate Gaussian prior that captures the within-cell variability. An expectation-maximization (EM) procedure alternates between penalized pixel-wise updates in latent space (E-step) and moment-based updates of the latent mean and covariance at the cell level (M-step). Per-channel amplitudes enter linearly and are eliminated by variable projection, so the optimization focuses on the non-linear physical parameters. After convergence, we map the latent parameters back into the constrained biological space to obtain per-pixel estimates, from which we compute per-cell means and standard deviations for the biological parameters summarized in Table 1.

We model each pixel independently with the same forward model described in Eq. (22), but use a vectorized form stacking the three time-resolved channels (D, S, and A) as one vector **y**_*p*_ ∈ ℝ^3N^, which takes the form

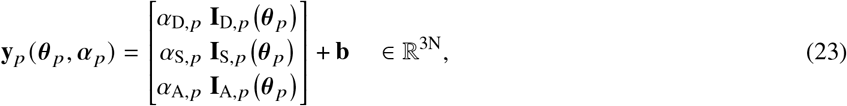

where **b** ∈ ℝ^3N^ is the vectorized form of **B** and represents the constant background. Throughout the paper, vectors/matrices that stack channels use the fixed order (D,S,A). Given the corresponding stacked measured signal from the three channels as **y**_m,*p*_ ∈ ℝ^3N^, the per-pixel maximum likelihood estimate (MLE, negative log-likelihood) is

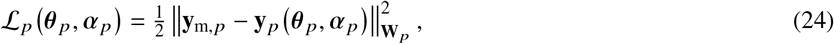

where **W**_*p*_ is the diagonal inverse-noise covariance for the stacked bins, and 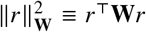. We pre-estimate the diagonal weights **W**_*p*_ from experimental calibration data using the procedure reported in our previous work (18). The per-channel amplitudes *α*_*p*_ enter linearly, so we eliminate them by variable projection. For fixed *θ*_*p*_,

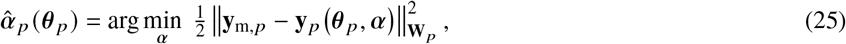

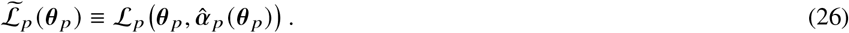

We solve this optimization for *θ*_*p*_ defined in Eq. (21), under the constraints in Eqs. (12) and (2). Our biological endpoint is the per-cell mean and standard deviation of each parameter across pixels

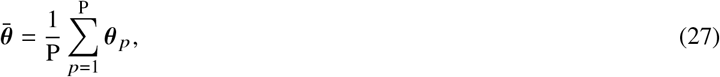

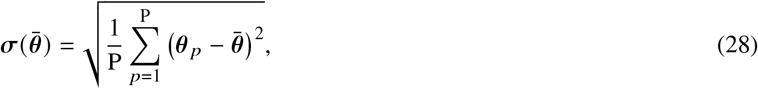

where P is the total number of pixels in the cell. We introduce an unconstrained latent parameter vector **z**_*p*_ ∈ ℝ^7^ for each pixel *p* = 1, …, P and use a smooth, invertible transformation to 𝒯 shuttle between the physical (constrained) space and the latent (unconstrained) space: **z**_*p*_ =𝒯 (*θ*_*p*_), *θ*_*p*_ = 𝒯^−1^ (**z**_*p*_). We chose the map 𝒯 so that the lifetime bounds and the probability simplex constraint for the fractions are automatically satisfied under the inverse map 𝒯^−1^, simplifying the within-cell pixel-to-pixel biological variability in the latent space. This allows us to place a simple i.i.d. Gaussian model across pixels in the latent space **z**_*p*_. Specifically, we treated the aggregate-associated lifetimes (*τ*_b_, *τ*_u_) as arising from multiplicative variability in the corresponding additional quenching rates (*k*_b_, *k*_u_), and we modeled these rates as approximately log-normal across pixels. This approach yields right-skewed lifetime variability while preserving the constraint *τ*_b_, *τ*_u_ ≤ *τ*_m_. For example, the lifetime transform takes the form 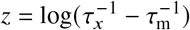, so that the inverse 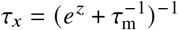 enforces *τ*_*x*_ ≤ *τ*_m_ for any *z* ∈ ℝ.

We modeled the acceptor-state fractions (*w*_bm_, *w*_um_, *w*_b_, *w*_u_) as a logistic-normal composition by placing a multivariate Gaussian model on log-ratio coordinates, which enforces positivity and the probability simplex constraint in Eq. (2) while capturing correlated trade-offs among classes. For example, given *j*, *k* ∈ {bm, ba, ua}, the fraction transform takes the form *z*_*j*_ = log (*w*_*j*_ /*w*_um_) with inverse 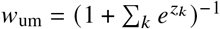 and 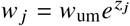 for these components. For the membrane densities *ρ*_m_, *ρ*_a_, we used the identity mapping in the latent representation, so that their coordinates enter 𝒯 unchanged. Thus, the transform 𝒯 is fully specified for all seven parameters, while no additional positivity or range constraint is imposed on the density components. In practice, the pooled latent Gaussian prior provided the stabilization needed for weakly identified pixels without introducing an explicit density transform. Consequently, we assume the pixel-wise latent vectors are i.i.d. and drawn from a shared multivariate Gaussian with mean ***μ***_*z*_ and covariance **R**_*z*_ ∈ ℝ^7×7^, which we initialize as diagonal. We thus have **z**_*p*_ ~ 𝒩 (***μ***_*z*_, **R**_*z*_).

To prevent boundary-seeking in weakly identified directions, we include an anchor prior on each pixel latent vector,

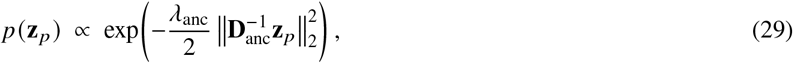

where **D**_anc_ = diag (**σ**_anc_) sets per-parameter scales and *λ*_anc_ is the regularization factor to control the strength of this prior so it does not bias the parameters. We used *λ*_anc_ = 0.1, and set **σ**_anc_ as 6 for the lifetimes, 10 for the densities, and 4 for the fractions.

This leads to the following iterative procedure.

**E-step (pixel-wise penalized estimate for z)**: Using Eq. (26) for the per-pixel negative log-likelihood, for fixed ***μ***_*z*_, **R**_*z*_, each pixel estimate is

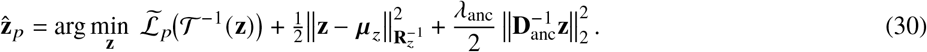

**M-step (pooled moments for z)**: With latent estimates 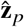, we update the mean and covariance as

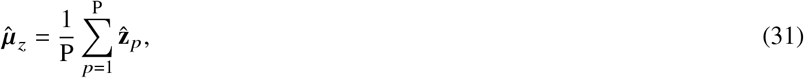

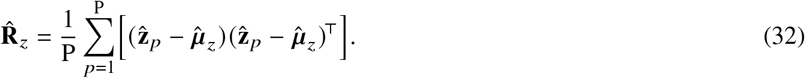

To stabilize the iterations, we apply mild damping using

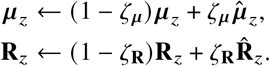

We set the damping values to *ζ*_***μ***_ = 0.2 and *ζ*_**R**_ = 0.1.

The E- and M-steps alternate until (***μ***_*z*_, **R**_*z*_) stabilize. We initialize EM with a uniform latent guess **z**^(0)^ corresponding to a generic physical parameter guess. In the first iteration, the pixel-wise optimization omits the pooled prior term in Eq. (30) and retains only the anchor regularization, while the physical constraints remain enforced through 𝒯^−1^. For iterations *i* ≥ 2, each pixel optimizer is warm-started using a convex combination of its previous latent estimate and a neighbor-averaged estimate,

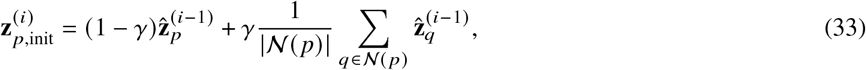

with *γ* ∈ [0, 1] controlling the strength of spatial conditioning and *p* containing the neighborhood of pixel *p*. In our results, we used *γ* = 0.5. We used neighbor averaging only to initialize the optimizer; it does not modify the objective in Eq. (30). This warm-start substantially improves stability and reduces solver iterations in low-SNR pixels. After convergence, we map the pixel estimates back to the physical domain via 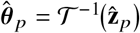 and compute per-cell summaries described in Eq. (27) and Eq. (28).

### 2.5 Monte Carlo data generation

We generated synthetic three-channel FLIM decays from the forward model in Eq. (22) using fixed calibration constants (Eq. (20)) and spatially varying ground-truth parameter maps *θ*_*p*_ (Eq. (21)) defined on an N_px_ × N_px_ grid and 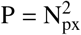 pixels. To impose within-cell spatial heterogeneity while retaining smooth parameter variation, we drew latent stationary Gaussian random fields (GRFs) using an FFT-based spectral method. Specifically, for each latent field we filtered spatial white noise in the Fourier domain with a squared-exponential power spectrum 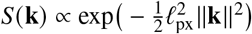, transformed back with an inverse FFT, and normalized to zero mean and unit variance. This construction yielded approximately stationary, smoothly varying fields with correlation length set by *ℓ*_px_ (here *ℓ*_px_ = 3).

We converted latent GRFs to physically constrained parameters using point-wise monotone transforms (using the 𝒯 transform defined earlier), with latent moments chosen to match the intended within-cell means and variances (reported in the results along with the simulation settings). Acceptor-state fractions, defined in Eq. (1) were generated as a spatial logistic-normal composition: we sampled three correlated latent “logit” GRFs relative to a reference component and mapped them to the probability simplex via a softmax transform, which produces strictly positive fractions that sum to one. We then calibrated the logit intercepts by a small fixed-point correction so that the spatial mean of the simulated fractions matched the target mean vector (51).

Aggregate-associated lifetimes (*τ*_u_, *τ*_b_) were generated from independent latent GRFs by first mapping each field through the standard normal CDF to obtain uniform variates, then applying inverse-CDF sampling for the desired marginal model. In particular, we enforced the upper bound in Eq. (12) by sampling a shifted log-normal distribution in the rate domain *k* = 1 /*τ* with a minimum rate *k*_min_ = 1/ *τ*_max_ (equivalently *τ* ≤ *τ*_max_), and transforming back to *τ*. This “Gaussian-field to uniform to target marginal” construction corresponds to a Gaussian-copula approach for prescribing marginals while inheriting dependence from a latent Gaussian model (52).

Membrane densities (*ρ*_m_, *ρ*_a_) were generated from latent GRFs and mapped to a bounded physical range (0, *ρ*_max_) using a smooth squashing transform; we report realized maxima in the Results section; we selected the latent mean and scale to place the simulated densities at the desired mean level with moderate spatial variation, and we report the realized mean and dispersion in the results.

For each pixel *p*, we computed the noise-free stacked decay **y**_*p*_ ∈ ℝ^3N^ from the forward model using the channel order (D,S,A) in Eq. (23). To emulate intensified wide-field FLIM noise, we added zero-mean Gaussian noise with bin-dependent variance proportional to the predicted signal. Let **y**_*p,c*_ denote the subvector for channel *c* ∈ {*D, S, A*}. We set

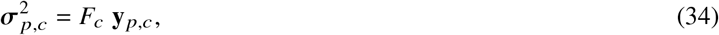

where *F*_*c*_ was chosen to enforce a prescribed peak SNR in each channel. We concatenated 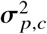 across channels to form 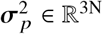 and generated noisy observations as

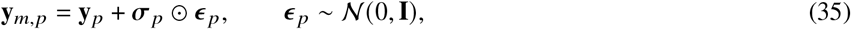

where ⊙ is the element-wise multiplication of the vectors, and ϵ_*p*_ is a standard normal random noise vector. The corresponding diagonal weights

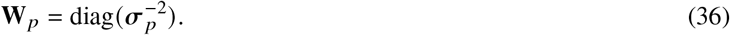

### 2.6 Calibration procedure

We estimated the fixed calibration constants in Eq. (20) from donor-only and acceptor-only control samples acquired with the same three-channel FLIM settings as the neuronal experiments. In each control dataset, we restricted analysis to high-signal pixels to avoid low-SNR bias and instability, using an intensity threshold on the peak gate in the dominant channel (Channel D for donor-only controls; Channel A for acceptor-only controls).

For each retained pixel, we fit a single-exponential decay model convolved with the measured instrument response function and coupled across the three channels through the pathway-mixing matrix in Eq. (18). The fit estimated the control lifetime (*τ*_d_ for donor-only; *τ*_m_ for acceptor-only) along with nuisance amplitude and background terms, and the relevant mixing coefficients. We used weighted nonlinear least squares (Gaussian MLE) consistent with Eq. (24).

Pooling pixel-level fits yielded *τ*_d_ = 3.9 ns for donor-only controls with *β*_a_ = 0.9 and *χ*_d_ = 0.2, and *τ*_m_ = 3.0 ns for acceptor-only controls. The acceptor-only mixing terms were negligible in our configuration, so we set *β*_d_ = 0 and *χ*_a_ = 0 for the experimental analyses. In the Monte Carlo simulations, we set all pathway-mixing terms to zero (i.e., *α*_d_ = *α*_a_ = *χ*_d_ = *χ*_a_ = 0) to isolate estimator performance under an idealized model without cross-talk. Because these cross-talk terms depend on the specific excitation/emission bands and optical configuration, their calibrated values are specific to our experimental design.

Other experimentally specified quantities, including the measured instrument response function and the background terms, are treated separately as known acquisition-model inputs and are not included in𝒩. Fixed geometric or photophysical quantities such as *R*_e_ and *R*_0_ are likewise prescribed independently from literature values rather than estimated from the control-sample calibration experiments, and are therefore not included in 𝒩.

### 2.7 Cell preparation

#### 2.7.1 Preparation of aSyn pre-formed fibrils

Purified human aSyn was concentrated to 5 mg/mL (347 *μ*M) using a 10 kDa molecular weight cutoff (MWCO) spin filter (Sartorius, Columbus, OH) operated at 4, 500 ×*g*. The concentrated protein solution (0.5 mL) was filtered through a 0.22-*μ*m syringe filter (Fisherbrand polyethersulfone, 33 mm) and incubated in a sterile 1.5-mL microcentrifuge tube at 37°C for 7 days with continuous shaking at 123 ×*g* in a Thermomixer (BT LabSystems BT917). The resulting fibrils were collected by centrifugation at 13, 000 ×*g* for 15 min and resuspended in 250 *μ*L of Dulbecco’s phosphate-buffered saline (DPBS; Cytiva). An aliquot of the fibril suspension was treated with guanidine hydrochloride (final concentration, 8 M) and incubated at 22°C for 1 h to dissociate fibrils into monomers. The aSyn concentration was measured via absorbance at 280 nm using a Nanodrop spectrophotometer, with an extinction coefficient of 5960 M^-1^cm^-1^. Fibrils were resuspended to a final concentration of 5 mg/mL, divided into 257 *μ*L aliquots, and stored at −80°C. Before use, fibril suspensions were sonicated in ethanol-sterilized tubes (Active Motif) using a cup horn sonicator (Qsonica Q700) set to 30% power (100 W/s) with a cycle of 3 s on and 2 s off, for a total “on” time of 8 min. Fibrils underwent 3 rounds of sonication. The bath temperature was maintained between 5°C and 15°C throughout the sonication.

#### 2.7.2 Preparation of adenoviral constructs

Adenoviral constructs encoding aSyn A53T-mVenus, Rab7-mT2, or mVenus were produced using the Virapower Adenoviral Expression System from Invitrogen (Carlsbad, CA). cDNAs encoding mVenus or mT2 were generated via PCR using the plasmids ecAT3.10 (Addgene #107215) and pRSETB-tdTomato-*ε*-CeN (53), both provided by Dr. Mathew Tantama (Wellesley College), as templates. cDNAs encoding human aSyn, and Rab7a were prepared via PCR using the plasmids pENTR-aSyn (54), GFP-Rab7 WT (Addgene #12605, deposited by Dr. Richard Pagano) as templates. GFP-Rab7 WT was provided by Dr. Robert Stahelin (Purdue University). PCR reactions were carried out using Phusion High-Fidelity DNA polymerase or Q5 Hot Start High-Fidelity DNA polymerase (NEB, Ipswich, MA). Overlapping PCR fragments encoding aSyn-mVenus, Rab7-mT2, and a single PCR fragment encoding mVenus, were subcloned into a variant of the entry vector pENTR1A carrying the human synapsin promoter (h-syn-P) (54), digested with KpnI and XhoI. The ligation reactions were carried out using the NEBuilder HiFi DNA Assembly kit (NEB). The pENTR1A-h-syn-P-aSynA53T-mVenus plasmid was generated by site-directed mutagenesis using overlap extension PCR to introduce the A53T mutation. The insert from each pENTR1A construct was transferred into the ‘promoter-less’ pAd/PL-DEST5 adenoviral expression vector (54) via Gateway recombination cloning (Invitrogen). The DNA sequence of each insert was verified by Sanger sequencing (Genewiz, South Plainfield, NJ). The resulting adenoviral constructs were packaged into virus via lipid-mediated transient transfection of the HEK 293A packaging cell line. Adenoviral titers were determined using the Adeno-X qPCR titration kit from Takara Bio USA (Mountain View, CA).

#### 2.7.3 Preparation and treatment of primary cortical cultures

Primary cortical cultures were prepared as described (55) by dissecting day 17 embryos obtained from pregnant Sprague-Dawley rats (Envigo, Indianapolis, IN) using methods approved by the Purdue Animal Care and Use Committee. Briefly, the cortical layers were isolated stereoscopically and dissociated by incubation with papain (20 U/mL) in sterile Hank’s Balanced Salt Solution (HBSS) at 37°C for 45 min. The dissociated cells were plated on poly-D-lysine-coated 8-well chambered glass plates (Cellvis) at a density of 75,000 cells/well in Neurobasal media supplemented with 2% (v/v) B-27 supplement, 5% (v/v) FBS, 1% (v/v) GlutaMAX, 50 U/mL penicillin, and 50 *μ*g/mL streptomycin. The next day, the plating media was replaced with Neurobasal media plus 2% (v/v) B-27 supplement, 1% (v/v) GlutaMAX, 10 U/mL penicillin, and 10 *μ*g/mL streptomycin. After 6 days in vitro (DIV = 6), the cultures were transduced with adenovirus encoding aSyn A53T-mVenus, Rab7-mT2, and/or mVenus under the control of the synapsin promoter at a multiplicity of infection (MOI) of 5. A subset of cultures were also treated with aSyn PFFs (final concentration, 6 *μ*g/mL). The cells were then incubated for 5 days, fixed with 4% (w/v) paraformaldehyde (PFA) in PBS for 15 min, and imaged via FLIM.

## 3 RESULTS

### 3.1 Simulation validation of hierarchical EM for three-channel cell-level readouts

To validate the estimator under controlled conditions, we analyzed synthetic three-channel datasets generated from the forward model using known parameters. A summary of the simulated values is provided in Table 2. We stacked the generated three-channel decays into a single vector as in Eq. (23). Then, we added signal-dependent Gaussian noise with SNR = 20 dB, defined here as the ratio of the peak bin mean to its standard deviation, to match the empirical noise level measured on our system. Figure 3 shows a representative noisy decay for a single pixel with the fitted prediction obtained by minimizing Eq. (24). The weighted residuals (bottom) are randomly distributed around zero, indicating that the model structure and noise weighting are consistent with the simulated data.

**Table 2:** Pixel-level bias and standard deviation (Std) within one representative simulated cell (400 pixels, SNR = 20 dB) for pixel-wise fitting (PW) and hierarchical EM (EM). Bias is computed as the pixel-average of 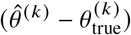, and Std is computed across pixels.

| Parameter | Simulated | Bias (PW) | Bias (EM) | Std (PW) | Std (EM) |
| --- | --- | --- | --- | --- | --- |
| $\tau_b$ (ns) | 1.70 | -0.40 | 0.11 | 0.91 | 0.21 |
| $\tau_u$ (ns) | 2.30 | -0.15 | -0.18 | 0.92 | 0.42 |
| $w_{bm}$ | 0.31 | -0.06 | -0.02 | 0.24 | 0.08 |
| $w_u$ | 0.31 | 0.07 | -0.04 | 0.31 | 0.07 |
| $w_b$ | 0.14 | 0.06 | 0.05 | 0.21 | 0.03 |
| $\rho_m$ ($10^{-3}$ nm$^{-2}$) | 5.00 | -2.39 | 0.16 | 8.72 | 1.94 |
| $\rho_a$ ($10^{-3}$ nm$^{-2}$) | 3.00 | 39.45 | -0.35 | 348.81 | 2.35 |

**Figure 3:**
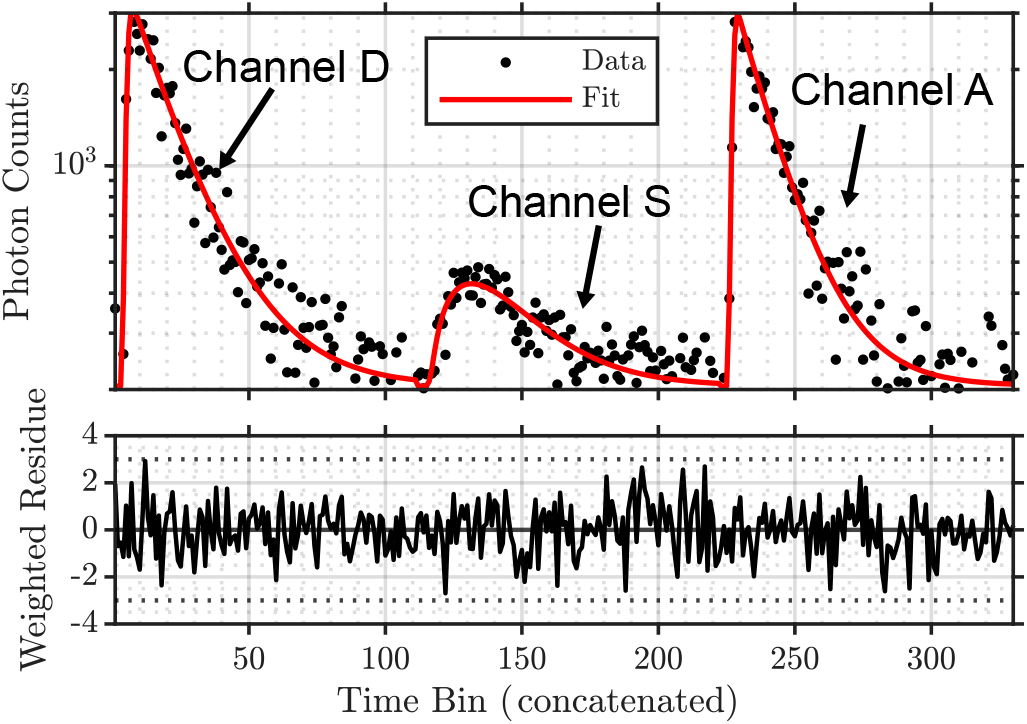
Representative time-resolved decay and fitted forward-model prediction for one pixel for the three-channel decay signals concatenated. Top: photon counts (markers) and fit (line) shown on a logarithmic photon-count axis; time bins from the D, S, and A channels are concatenated along the horizontal axis. Bottom: weighted residuals for the same pixel.

Next, we quantified the accuracy of cell-level biological readouts. We compared the proposed hierarchical EM framework, using 5 iterations, against a standard pixel-wise approach, where parameters are fitted individually per pixel using Eq. (24) and then averaged over the cell. Figure 4 summarizes the distribution of estimated parameters across five independent realizations, each with P = 400 pixels. The hierarchical EM estimates (red, right) exhibited markedly lower bias and variance compared to the pixel-wise estimates (blue, left), with distributions that are tightly centered around the simulated values (black lines). This reduction in bias and variance is evident for most parameters.

**Figure 4:**
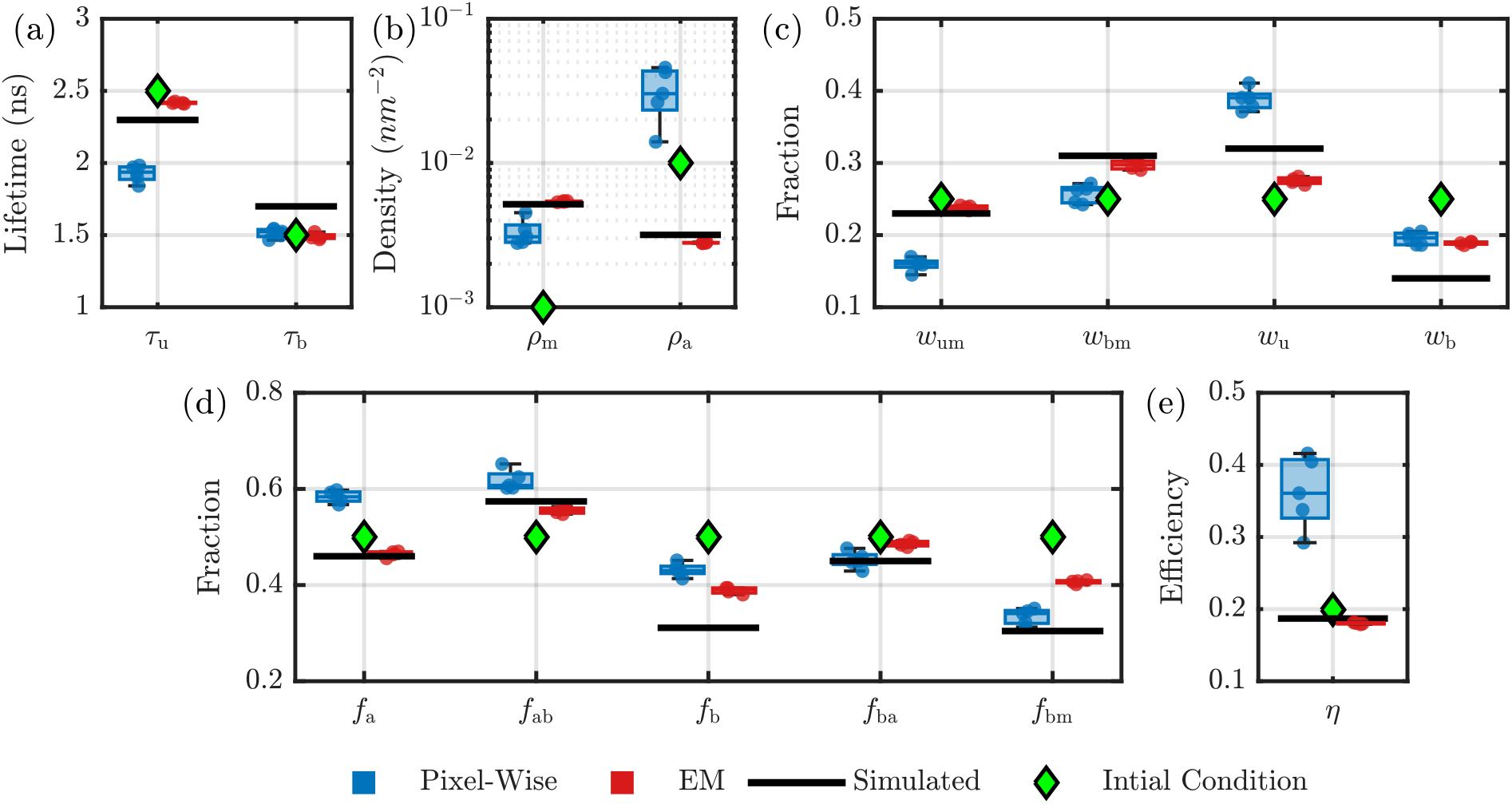
Cell-level parameter estimates across the Monte Carlo ensemble (5 repetitions, *P* = 400 pixels per cell, SNR = 20 dB). Boxplots summarize per-cell means computed using Eq. (27), obtained from pixel-wise fitting followed by averaging and from hierarchical EM algorithm (EM). Black horizontal lines indicate the simulated values; green diamonds indicate initialization. Panels report: (a) lifetimes (*τ*_u_, *τ*_b_), (b) surface densities (*ρ*_m_, *ρ*_a_), (c) acceptor-state fractions (*w*_um_, *w*_bm_, *w*_u_, *w*_b_), (d) derived biological readouts (Table 1), and (e) donor-acceptor FRET efficiency (*η*).

The numerical stability of the EM updates is illustrated in Fig. 5, which shows the evolution of cell-level parameter estimates over EM iterations. The shaded band denotes ±1 standard deviation across pixels within the cell at each iteration. Most parameters converged rapidly and stabilized within ~5 EM iterations. Beyond this point, the within-cell standard deviation across pixels became tightly constrained, reflecting the regularization of individual pixel estimates toward the population mean. However, the membrane-bound aggregate fraction converged more slowly.

**Figure 5:**
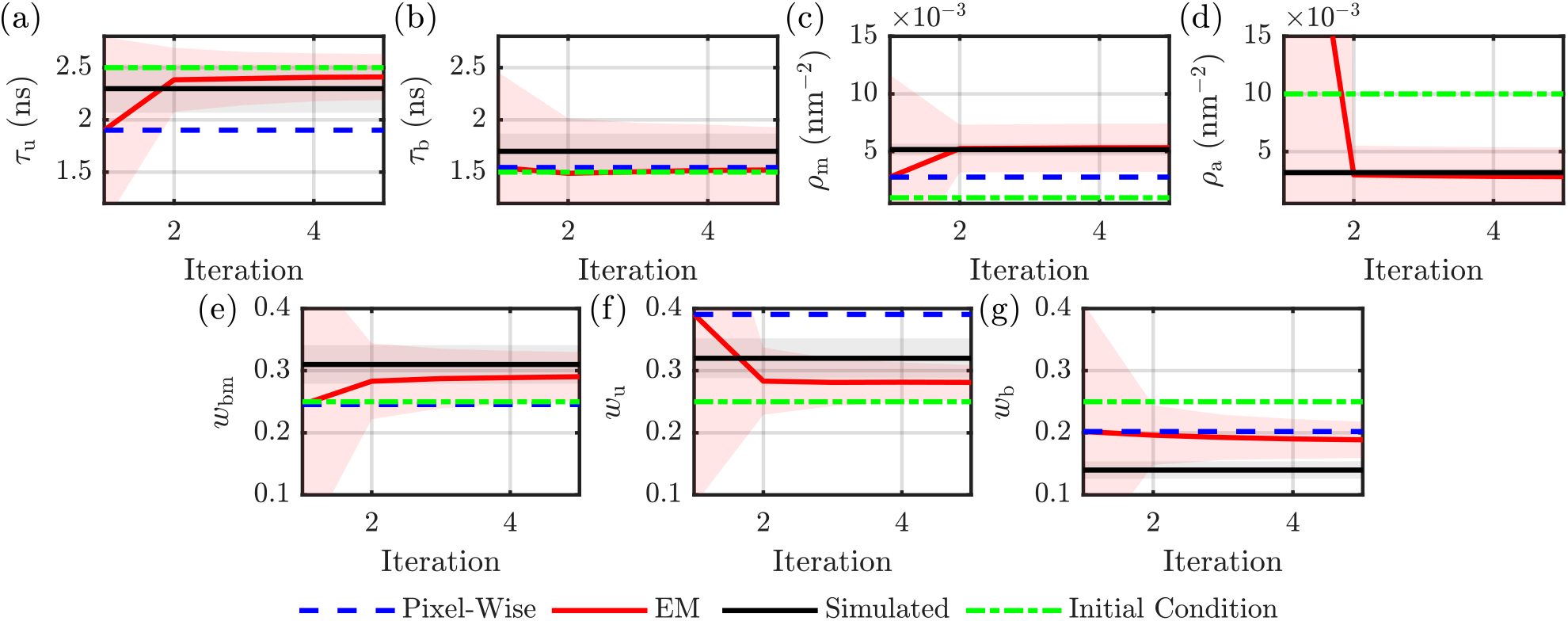
Parameter trajectories versus EM iteration for one simulated cell. Dashed blue: pixel-wise estimate shown as a fixed reference; solid red: hierarchical EM estimate (EM); black: simulated values; green: initialization. The shaded band denotes ±1 within-cell standard deviation across pixels at each iteration. Panels report: (a) membrane-unbound aggregate lifetime (*τ*_u_), (b) membrane-bound aggregate lifetime (*τ*_b_), (c) monomer surface density (*ρ*_m_), (d) aggregate surface density (*ρ*_a_), (e,f,g) acceptor-state fractions (*w*_bm_, *w*_u_, *w*_b_)

Next, we assessed the effect of hierarchical EM on pixel-level estimates by evaluating the spatial fidelity in Fig. 6 for one of the parameters, namely the lifetime of membrane-unbound aggregates (*τ*_u_). Figure 6(a) shows the simulated *τ*_u_ map. Figure 6(b) shows the pixel-wise estimates, and Fig. 6(c) shows the hierarchical EM estimates. The pixel-wise estimation produced maps corrupted by high-spatial-frequency noise. In contrast, the hierarchical EM recovered the smoother spatial structure of the simulated values. The improvement map, shown in Fig. 6(d), confirms that estimation error is reduced for most pixels, demonstrating that the method improves per-cell quantification while preserving the large-scale spatial structure present in the simulated field.

**Figure 6:**
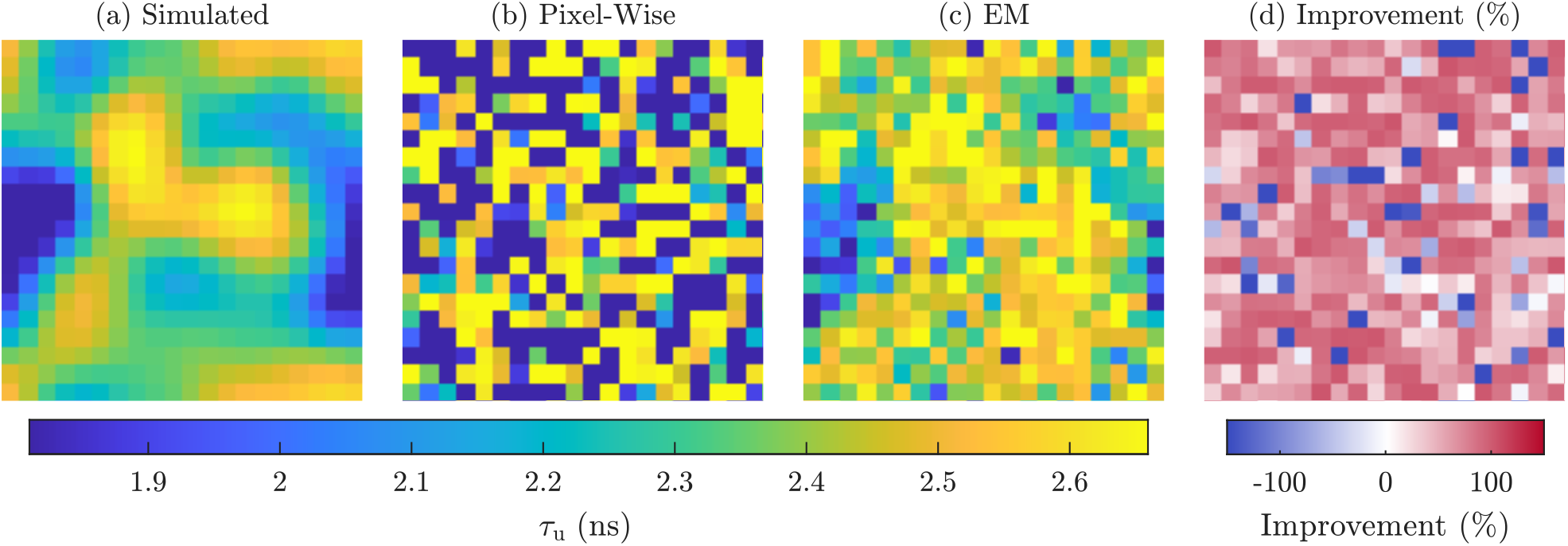
Pixel-level maps for *τ*_u_: (a) simulated value, (b) pixel-wise estimates (PW),(c) hierarchical EM estimates (EM), and(d) percent improvement computed using 100 (Err^PW^ −Err^EM^)/*θ*, where 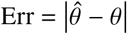. Positive improvement indicates reduced absolute error under EM relative to PW.

Finally, Table 2 provides a quantitative comparison of pixel-level estimator performance. The hierarchical approach substantially reduced the standard deviation of the estimates; for example, the standard deviation for the aggregate-associated lifetime *τ*_b_ was reduced by approximately 77%, and the bound-aggregate fraction *w*_b_ by approximately 84%. While bias varied by parameter, the hierarchical model generally maintained or improved accuracy, particularly for the critical density parameter *ρ*_m_ and acceptor fractions.

### 3.2 Fixed-cell neuronal validation with aSyn PFF seeding

To demonstrate performance on experimental data and to probe biologically relevant changes in aggregation, we applied the hierarchical three-channel analysis to fixed neurons expressing Rab7-mTurquoise2 (Rab7-mT2) and aSyn-mVenus under two experimental conditions: cells seeded with exogenous aSyn pre-formed fibrils (+PFF) and unseeded controls (−PFF). For each condition, regions of interest were segmented and analyzed jointly across the three channels using the forward model and hierarchical EM estimator.

Figure 7 shows box plots of the estimated parameters and the corresponding biological readouts (Table 1) across cells for +PFF (blue, left) and −PFF (red, right), with calibration parameters held fixed as described in the Methods. Notably, both membrane-bound and membrane-unbound aggregate lifetimes *τ*_u_ and *τ*_b_ were systematically reduced, with significant differences by one-way ANOVA, in the +PFF condition relative to controls, consistent with increased FRET and enhanced aSyn aggregation at or near the membranes. These results were based on a total of 18 cells (*n*_+*PFF*_ = 9, *n*_−*PFF*_ = 9) sampled from 2 animals prepared according to the procedure described in the Methods. Several aggregation-related fractions also shifted in the expected direction in +PFF-treated cells, indicating a higher aggregate burden and altered membrane-proximal aggregate fractions. These results demonstrate that the proposed three-channel hierarchical analysis can resolve PFF-induced changes in aSyn aggregation state at the per-cell level, even when per-pixel noise would obscure such differences in conventional pixel-wise maps.

**Figure 7:**
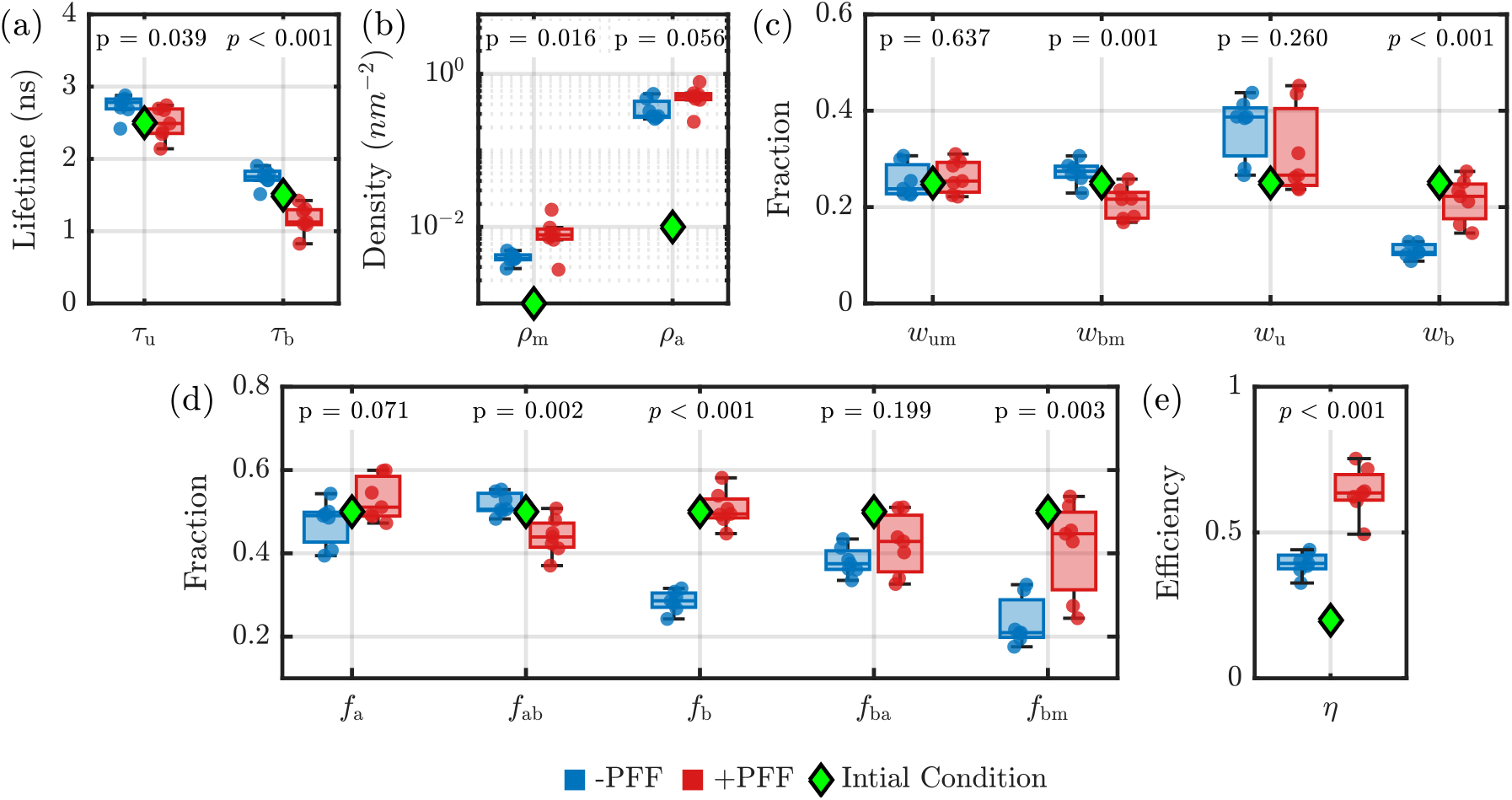
Cell-level parameter estimates for fixed neurons expressing Rab7-mTurquoise2 and aSyn-mVenus under alpha- synuclein pre-formed fibril seeding (+PFF) and unseeded controls (−PFF). Box plots summarize per-cell means obtained with the hierarchical EM estimator; green diamonds indicate initialization. Colors denote condition (+PFF: blue, left; −PFF: red, right). Group differences were tested by cell-level ANOVA. These data were collected from 18 cells total (*n*_+*PFF*_ = 9, *n*_−*PFF*_ = 9) sampled from 2 animals prepared according to the procedure described in the Methods. Panels report: (a) lifetimes (*τ*_u_, *τ*_b_), (b) surface densities (*ρ*_m_, *ρ*_a_), (c) acceptor-state fractions (*w*_um_, *w*_bm_, *w*_u_, *w*_b_), (d) derived biological readouts (Table 1), and (e) donor-acceptor FRET efficiency (*η*). Results showed shifts consistent with increased aggregation and increased membrane-proximal interactions in +PFF-treated neurons.

## 4 DISCUSSION

This work combines three-channel FLIM-FRET with a forward model and hierarchical inference to quantify membrane-proximal aSyn aggregation in neurons at the cell-level. The approach targets a central challenge in live-cell aggregation studies, where multiple molecular classes contribute photons within a diffraction-limited voxel and conventional pixel-wise fits become unstable under realistic photon budgets (17). We validated the method experimentally. In the +PFF condition, the analysis detected systematic reductions in both membrane-bound and membrane-unbound aggregate-associated lifetimes (*τ*_b_, *τ*_u_), alongside shifts in aggregation-related fractions, which aligned with increased aggregation burden and altered membrane-proximal aggregation in seeded neurons (30, 32, 34). These results support the view that membrane interactions can shape early aggregation pathways (5, 27, 28). A primary benefit of the three-channel measurement is improved identifiability of the latent mixture states relative to single-channel FLIM-FRET. Channel D constrains donor quenching produced by donor-acceptor proximity at the membrane. This is quantified using a membrane-FRET model that accounts for density-dependent transfer (39–41) while enforcing steric exclusion limits between the fluorescent protein structures (44). Channel S adds the sensitized-emission kinetics that couple donor decay to acceptor excitation, which reduces degeneracy between amplitude mixing, donor lifetime shortening, and spectral cross-talk that often limits single-channel inference (48–50). Channel A independently constrains acceptor lifetime shortening associated with dense microenvironments and self-quenching, which provides an orthogonal signature of aggregation (11, 12). Jointly fitting these channels under shared physical parameters therefore separates membrane proximity effects from aggregate-associated quenching and yields interpretable, condition-comparable readouts that do not require optical super-resolution (14–16).

The hierarchical EM estimator improves cell-level quantification by pooling information across pixels in a statistically principled way, but it does not collapse the data into a single decay per cell. Conventional global analysis gains stability by fitting one decay per cell, yet it necessarily discards within-cell spatial heterogeneity (19–21). In contrast, the hierarchical formulation still estimates per-pixel parameters and retains spatial structure, while shared cell-level latent statistics stabilize low-SNR pixels. Segmentation-based global fitting partially bridges this gap by grouping pixels into regions before fitting (56), but it still suppresses variability within each segment and makes results depend on segmentation choices. Fit-free phasor analysis offers a complementary, model-agnostic way to characterize lifetime and FRET behavior (57), although it does not directly yield the same physically parameterized mixture fractions and membrane-density readouts targeted here. Pixel-wise fitting treats each pixel as an independent inverse problem, so weakly identified parameters can drift toward constraint boundaries and averaging can then produce biased and high-variance cell summaries (17). The hierarchical model mitigates this failure mode by coupling pixel parameters through an unconstrained latent Gaussian representation and updating cell-level moments during EM (35, 36). This coupling regularizes noisy pixels toward cell-consistent solutions while still allowing within-cell variability through the learned covariance, which explains the reduced bias and variance observed in Monte Carlo validation and the cleaner spatial maps that can be tuned further via the prior strength, damping, and spatial conditioning settings.

Several possible limitations arise from the model structure and calibration choices. First, the membrane-FRET component assumes acceptors are randomly and independently distributed in the membrane plane and uses a Wolber-Hudson type approximation, therefore deviations from these assumptions, such as non-Poisson clustering, curvature, or heterogeneous membrane geometry, could introduce model mismatch (39–41). Second, the analysis compresses membrane heterogeneity into two surface densities (*ρ*_m_, *ρ*_a_), which may not represent a continuum of local densities or additional organizational states in neurons. Consistent with this limitation, quantitative neuronal imaging has reported strongly heterogeneous aSyn membrane association, including diffuse pools coexisting with bright puncta and broad per-vesicle copy-number distributions (58). In addition, aSyn can form synaptic multimers that concentrate within vesicle clusters and restrict vesicle motility (59). The third limitation is that the inference depends on the marker topology: Rab-family small GTPases are prenylated and require membrane insertion for function, while GDIs sequester inactive Rab in the cytosol by masking prenyl groups (60). If the fluorescent Rab marker (or a fraction of the acceptor population) instead behaves as a luminal label or reports intraluminal vesicles, then donor-acceptor separations no longer follow an in-plane 2D geometry. The effective separation increases by approximately the bilayer thickness plus linker lengths, which can suppress FRET and bias inferred densities. In that scenario, the forward model must explicitly account for opposite-leaflet or volumetric acceptor distributions, and the experiment should include topology controls for the chosen membrane fluorophore. Fourth, the estimator holds calibration terms fixed (unquenched donor and acceptor lifetimes, bleed-through, cross-excitation, and the closest-approach distance *R*_e_), so calibration errors could propagate into inferred physical parameters through the pathway mixing matrix and the coupled three-channel model (48–50). If needed, selected terms can be treated as additional variables with informative Bayesian priors, which permit small, physically plausible deviations while maintaining identifiability. The practical benefit depends on the experimental configuration and the magnitude and uncertainty of the calibration terms. These limitations motivate sensitivity analyses to calibration perturbations and targeted model extensions when the data indicate systematic residual structure.

The framework should generalize to other membrane-mediated assembly processes where a proximity reporter isolates membrane-proximal interactions and a second signature reports biochemical state. In particular, any system where donor lifetime reports nanoscale proximity and acceptor lifetime reflects local microenvironment or self-quenching can benefit from three-channel constraints and pooled inference, including other protein-misfolding systems and membrane-proximal condensate or clustering phenomena. Lifetime-based FRET readouts have quantified state-dependent organization and assembly in other membrane systems, including EGFR organization and K-Ras nanocluster formation (61, 62).

Looking forward, live time-lapse studies can test whether the inferred per-cell metrics track aggregation trajectories and respond predictably to perturbations such as lipid composition changes, trafficking modulation, or genetic variants that alter membrane affinity (38). Parameter-robustness studies, including calibration sensitivity and alternative noise likelihoods, can further strengthen interpretability and support broader deployment across microscopes and labeling strategies.

## 5 DATA AVAILABILITY

Data are available upon request.

## 6 AUTHOR CONTRIBUTIONS

AS designed the method, ran the simulations, performed the experiments, and wrote the manuscript. WQ prepared the neurons. CR and KW coordinated the project in their respective laboratories and revised the manuscript.

## 7 DECLARATION OF INTERESTS

The authors declare no conflict of interest.

## 8 ACKNOWLEDGMENTS

This work was supported in part by the National Science Foundation (CBET 1937986 and 2330643, and CCF 1909660), the National Institutes of Health (R21NS105048 and R21NS135424), and by the Michael J. Fox Foundation.

## Notes

### Competing Interest Statement

The authors have declared no competing interest.

